# Predicting gene expression responses to environment in *Arabidopsis thaliana* using natural variation in DNA sequence

**DOI:** 10.1101/2024.04.25.591174

**Authors:** Margarita Takou, Emily S Bellis, Jesse R Lasky

**Affiliations:** Pennsylvania State University, University Park, 16802, USA; Department of Computer Science, Arkansas State University, Jonesboro AR

**Author notes:** **Corresponding author:** Margarita Takou. **email addresses:** Emily S Bellis, Jesse R Lasky.

**Keywords:** low frequency variants, gene expression prediction, machine learning, regulatory elements, evolution, genotype to phenotype map

## Abstract

The evolution of gene expression responses is a critical component of adaptation to variable environments. Predicting how DNA sequence influences expression is challenging because the genotype to phenotype map is not well resolved for *cis* regulatory elements, transcription factor binding, regulatory interactions, and epigenetic features, not to mention how these factors respond to environment. We tested if flexible machine learning models could learn some of the underlying *cis-*regulatory genotype to phenotype map. We tested this approach using cold-responsive transcriptome profiles in 5 diverse *Arabidopsis thaliana* accessions. We first tested for evidence that *cis* regulation plays a role in environmental response, finding 14 and 15 motifs that were significantly enriched within the up- and down-stream regions of cold-responsive differentially regulated genes (DEGs). We next applied convolutional neural networks (CNNs), which learn *de novo cis-*regulatory motifs in DNA sequences to predict expression response to environment. We found that CNNs predicted differential expression with moderate accuracy, with evidence that predictions were hindered by biological complexity of regulation and the large potential regulatory code. Overall, approaches to predict DEGs between specific environments based only on proximate DNA sequences require further development, and additional information may be required.

## Introduction

Adaptation is often a complex process that occurs via selection on traits and the underlying genetic mechanisms (1). The genetic basis of ecologically important quantitative traits often involves mutations with effects on gene expression. In variable environments, the evolution of gene expression responses to environment (expression plasticity) are likely a critical component of adaptation (2). These processes are often studied using evidence from variation in DNA sequence and mRNA abundance. For instance, the incorporation of transcriptomic information into genome wide association studies (GWAS) can help identify genes controlling a specific phenotype (3). Association or linkage mapping approaches can be used to map expression quantitative trait loci (eQTL), and allele-specific expression can determine the presence of *cis* or *trans* acting eQTL (4–6). However, the integration of sequence and expression data in evolutionary quantitative genomics is hindered by gaps in our understanding of how DNA sequence influences expression. In particular, understanding the genetic basis of expression plasticity might be beneficial to applied biology given the importance of predicting organismal responses to environmental changes(Keagy et al. 2023).

Gene expression responses to the environment differ among genes in the genome and among different natural genotypes due to DNA sequence variation. Differences in gene expression among genotypes may be driven by sequence differences at other loci in the genome, which regulate expression of a given gene (*trans* effects). At the same time, differences among genes and genotypes may arise from differences in sequence proximal to the gene that determine how its expression is regulated (*cis* effects). Researchers have documented cases where mutations in genes involved in environmental sensing (8,9) or in environmentally responsive genes that bind DNA sequences (10,11) have downstream effects on expression of other genes. Additionally, researchers have characterized how some *cis* mutations influence DNA binding by environmentally responsive transcription factors, ultimately altering environmental responsive expression (12–14). Transcription factors that tend to show expression responses to environment have been documented, and the DNA sequence motifs they bind to have been roughly characterized (15). However, across the genome there are many transcription factors that often respond to a single environmental stressor and interact, and this knowledge is currently too limited to predict how genetic variation in DNA sequences influence genome-wide expression responses to environment.

The large number of *cis*-regulatory motifs allows the investigation of genomic and evolutionary properties, which are associated with their distribution across the genome (16–18). Genome-wide expression profiling in association mapping or linkage mapping populations as well as allele-specific expression studies have revealed that *cis* expression quantitative trait loci (*cis-*eQTL) are abundantly segregating within species. *Cis-*regulatory mutations have been of particular importance for evolutionary biology because some have suggested they may be less likely to be deleterious than mutations in transcription factors, which are expected to be more pleiotropic (19,20). Relatedly, the possibility of easier dissection of the genotype to phenotype map for *cis-*eQTL (versus *trans-*acting mutations likely influencing multiple genes’ expression) combined with the abundance of *cis-*eQTL may be an opportunity to build a more integrative understanding of quantitative trait genetics.

Using machine learning methods in fields such as genomics and population genetics can help navigate increasingly bigger datasets and reveal complex patterns (21). Especially in gene regulation and evolution, deep learning approaches can have an advantage over traditional methods in decoding enhancer grammar of gene regulation, as these models can learn complex *cis*-regulatory rules in a precise manner not biased by current knowledge (22) and can outperform more traditional methods, like clustering by kmer or linear regression in predicting gene expression (23). For example, training a machine learning model on the imputed *cis* haplotypic information of RNA expression, Giri et al. (2021) managed to exclude the impact of trans effects on gene expression variation in maize. The prediction was less variable than those using SNPs and had an increased accuracy for predicting within population variance (24). Random forest models successfully classified protein coding vs non-protein coding genes in maize (25). Moore et al. (2021) showed how known regulatory sites of Arabidopsis thaliana and transcription factors (TFs) interactions can be used in machine learning to predict clusters of gene co-expression related to stress response (26). Similarly, in Arabidopsis thaliana, information on TF binding sites has been harnessed to predict differential gene expression under stress such as response to Fe (27), high salinity (28) and combined heat and drought stress (29). However, in all of these cases, the authors used additional -omics information, such as chromatin accessibility, or co-expression clusters, during deep learning. There is increasing evidence that genetic diversity on the *cis* elements of plants is under selection (30). Deep learning algorithms could be used for genome wide identification of deleterious and adaptive mutations, which is a prerequisite for the genetic improvement of crops (31). Using upstream and downstream genetic coding information could improve the prediction of the role of *cis* regulatory elements in differential gene expression, in contrast to using only upstream regions in maize (32), especially when more than one genotype was included in the analysis (33). However, those methods do not capture distal regulatory elements well if at all (34) and the source of variability can impact their efficiency (35).

In *Arabidopsis thaliana,* whose importance in studying evolution and ecology patterns is growing over the years (36), the conserved noncoding sequences are mostly concentrated close to the coding region and impact gene regulation (37), as *cis*-regulatory variants are usually in linkage with the expressed transcript (16,38,39). Even though there are conserved gene reactions between species and populations, the increased genetic diversity within populations or species causes phenotypic variance. Genes, whose expression is affected by genotype x environment interactions (GxE), tend to show allele frequency correlations with climate and have elevated genetic variation in stress responsive transcription factor binding sites (40). Known TF binding site motifs of *A. thaliana* have been used to identify putative binding sites and then train a deep model for predicting *cis* changes in tomato (41).

Our aim was to use genetic variation among genes within individuals and among genotypes in the regions upstream and downstream of genes to predict expression response to environment in *A. thaliana.* We aimed to predict in a both a more general sense than other studies by using the sequences instead of known TFBS, and in a more restrictive sense by classifying gene expression in specific environments within one species. For the analysis we used a published dataset on differentially expressed genes (DEGs) during cold acclimation of diverse *A. thaliana* genotypes (42). Response to cold stress not only has some well characterized signaling pathways, transcription factors and transcription factor binding sites, but also has been connected to adaptation to local environments (43–45). Hannah et al. (2006) had identified approximately 1,500 up regulated DEGs and down regulated DEGs out of around 20,000 genes for multiple accessions as a response to cold acclimation and stress (S1 Table). Here, we focus on the five *A. thaliana* accessions *Can*, *Col-0*, *Cvi*, *Ler* and *Rsch*, for which the pseudogenomes from whole genome resequencing data are published (46). We included multiple genotypes in the analysis, so as to capture some of the natural genetic diversity and potential GxE responses.

We first tested whether there are specific sequence motifs enriched along those regions of the differentially expressed genes that would suggest *cis* regulation drives some aspect of cold expression response. We determined if these motifs were related to known stress-responsive transcription factors (TFs). Then, we tested whether several naïve deep learning models could predict expression response, without a priori designation of important motifs, by using only the sequences upstream and downstream of genes.

## Material and Methods

### Transcriptome and genome data

We used the publicly available dataset of differentially gene expression during cold acclimation by (42). In brief, the authors used multiple accessions of *A. thaliana* to test their cold acclimation potential. They found that approximately 4,000 genes respond to cold acclimation, indicating regulatory pathways being involved in their control. Out of the accessions used in the original study, we used the five accessions, *Col-0, Ler, Cvi, Can* and *Rsch* (S1 Table) whose pseudogenomes are available in https://1001genomes.org/. For comparison with a potentially more simple transcriptomic change, we used the differential gene expression of 1,280 genes in the *Col-0* background of a *tt8* knock out line. The *tt8* gene regulates biosynthesis of anthocyanins, which we reasoned would be a narrower regulatory network than that involved in environmental responses (47).

### Searching for sequence motifs using STREME

Transcription factors bind DNA in a sequence specific manner, but not all sequences are known nor is the exact relationship between sequence and binding affinity known. For this reason we began with a *de novo* search for sequence motifs in the upstream and downstream regions of the coding sequence, as those often contain such signals. We used samtools faidx v1.10 (48) to extract the 1kbp upstream and downstream regions of each gene present in the dataset, including the first and last part of the coding region as TFBS often bind there. Specifically, we extracted the 500bp upstream and 500bp downstream of the gene’s TSS for the upstream regulatory coding region, and the 500bp upstream and 500bp downstream of the gene’s end transcriptional site for the downstream regulatory coding region. We then used the software STREME (49) to calculate significantly enriched ungapped motifs within the set of up- and down-regulated DEGs in relation to their presents in non-DEGs. For both groups we used as a control a randomly down-sampled set of the non-DEGs and stopped the search for motifs when we failed to identify motifs with high enough confidence (e-value > 0.05) three times in a row. This way, the statistical significance of each motif was determined based on whether it can classify sequences as DEGs or non-DEGs.

Because sequence motifs may have a variety of causes and functions, and not all are likely TFBS, we next sought to determine whether motifs discovered *de novo* by STREME specifically correspond to known transcription factor binding sites. We searched for significant overlaps of at least 5bp between the discovered enriched motifs and the JASPAR 2022 CORE (50) plant transcription factor binding sites database with MEME suite’s online tool TOMTOM (51). Significant overlap was determined by scoring the match of all possible alignments between sequence and known motif and summing those scores to produce p values, q values, and e values.

### Random forest analysis

To determine whether known transcription factor binding sites regulate the response to cold, we attempted to predict up regulated or down regulated DEGs vs non DEGs based on their presence or absence in the regulatory regions. We used a dataset (52) of TF putative binding sites 1000 bp upstream of each gene, which were predicted from *A. thaliana* using JASPAR2018 Bioconductor package (JASPAR) and then matched to the aligned regions of *A. thaliana*. We treated the count of each TF binding site per gene as a separate feature and additionally determined the total number of TF binding sites per gene. Taken together, these descriptors of genomic variation provided a total of 413 features, of which we could assess the predictive impact on gene regulation.

We trained a random forest model for *Col-0* using the ranger library in R (54) with up-regulated DEGs in response to cold and non-DEGs as variable and all compiled features of gene regulation as predictor variables. We set the number of trees of the forest to 500, the number of variables to possibly split at each node (mtry) to 200, and the impurity mode to impurity_corrected. From the trained random forest, we extracted the relative importance of each predictor variable as well as the out-of-bag prediction error, which is informative about the predictive power of the model. Finally, we estimated a *p* value for each predictor variable using the “Altmann” method with 100 permutations. We repeated the random forest analysis for all down-regulated DEGs. We estimated the prAUC in R library using the function prauc and extracted its significance by permuting 100 times the gene identity for each set of variables.

### Training Convolutional Neural Networks

We employed a more naive machine learning approach to determine whether we can predict DEG using regulatory regions. We trained convolutional neural networks (CNNs) to identify a model that would predict whether each gene was differentially expressed in the stress conditions relative to the control with high accuracy. We prepared the dataset by separating the genes into a training and test set for each accession, incorporating information about the gene families that they belonged to (55). Because sequences may carry spurious associations with expression that are due to shared evolutionary history, we followed the method of Washburn et al. (2019) and split genes into gene families and then tested predictions in gene families not used for training.

In the end, we used 68,979 (80%) genes, from which the 20% (13,796 genes) was always retained as the validation set, and 17,475 (20%) as the test set out of the total 86,454 genes across all five genotypes. Each unique gene could be present multiple times in the dataset, representing each accession. The proportion of up- and down-regulated DEGs to the non DEGs was approximately 1:11 in both sets (S1 Table). For each gene, we used samtools faidx v1.10 (48) to extract the 1.5kbp upstream and downstream regions of each gene from the published pseudogenomes, using custom scripts. Specifically, we extracted the 1,000 upstream and 500 bp downstream of the gene’s transcriptional starting site (TSS) for the upstream regulatory coding region, and the 500 bp upstream and 1,000 bp downstream of the gene’s transcriptional ending site for the downstream regulatory coding region. Those sequences were converted into one hot encoding of each nucleotide in python v3.6, to be used as input for the training and testing of the CNN models. We then created the labels needed for binary classification of DEG during training by scoring the up-regulated DEGs as 0 and the non-DEGs as 1. As the number of up-regulated DEGs was much smaller than non-DEGs, we oversampled with replacement the up-regulated DEGs to a ratio of 1:1 between the two groups. This process was repeated for training models to classify genes as down-regulated DEGs and non-DEGs.

We used the prepared inputs and labels of the training set to perform a grid search of tuning parameters, as we wanted to identify both the models with the best prediction of our dataset, as well as how different tuning parameters influence model accuracy using the regulatory regions of multiple accessions. The grid search was performed on a total of 1,344 combinations of hyperparameters (55) by training each model for 50 epochs using the python library tensorflow-gpu v2 (56). The tuning parameters included: the number of convolutional layers, the filter of each convolutional layer, the width of each convolutional layer, the pool width and stride, the number of dense layers and their units, as well as the dropout rate. A list of the specific values used to create all the different combinations is provided in S2 Table. During each training phase, we used the prAUC values of the validation set to tune, select and save the best trained model for further evaluation. For each trained model, we extracted the loss, accuracy, and the probability-based accuracy under the curve (prAUC) of the training, validation and testing of each model to compare the accuracy of the models. Moreover, we estimated the proportion of correct predictions of each class within the test set so that we could assess the potential overfitting of each model to predicting one class or the other. We estimated this within class accuracy as the number of times that the class was predicted correctly divided by the total number of genes belonging in this class. In this context, class refers to up-regulated or non-DEG genes. The absolute difference of within class accuracies, referred to in the text as prediction accuracy between the two classes, was also estimated. We repeated this process for all down-regulated DEGs.

### Determining factors influencing accuracy of CNNs

To determine the degree to which predictions relied on upstream sequence versus both up and downstream sequences, we repeated the above analysis described for the regulatory regions of all accessions by using only the upstream regions.

Because genetic variation among genotypes in expression can be driven by *trans-*effects (from other loci far from the gene), we sought to control for some of this variation. In particular, we focused on known loss of function variation at large effect CBF transcription factors that are involved in adaptation to different temperatures, first by training and evaluating CNN models on those lines (*Can* and *Cvi),* which have all 3 CBFs functional copies differentially expressed between the experimental conditions (45,57) and also in those lines (*Rsch, Col-0* and *Ler*), which have no intronic or other SNP differences. The CBFs, or *C-REPEAT BINDING FACTORs* are a set of 3 transcription factors that regulate cold acclimation and segregate for loss of function variation at high frequency in *Arabidopsis thaliana*, with loss of function variants associated with warmer climates (45,58). Hannah et al. (2006) previously showed with our dataset that intact CBF function explained a substantial portion of variation in cold response transcriptomes and Des Marais et al. (2017) showed that genetically variable cold responses were enriched in central locations of coexpression networks.

To determine the role that genetic variation among genotypes, versus only among genes within a genotype, we trained models with only *Col-0* genes as the set. We repeated the training while also oversampling the dataset to be the same size as for all five accessions combined to control for underpowered training due to the smaller dataset.

To assess whether models may have been hindered by factors such as model architecture or data complexity we introduced a “spike” of 5bp in the upstream region of up-regulated DEGs. This way we created a positive control that can help validate the capability of the models to recognize the up regulated DEG group. We replaced the first 5bp of the up-regulated DEGs with the sequence AAGGG. Then, we evaluated each trained model in two dimensions; both based on their accuracy of predicting the evaluated set, but also based on their accuracy when predicting non spiked data.

### Evaluating and interpreting model training

We compared the area under the precision-recall curve (prAUC), cluster accuracy, as well as the difference of the two classes accuracy with a Kolmogorov Smirnov test using the function ‘ks.test’ in R v4.1.2. To see which hyperparameters had a significant influence on those CNN performance metrics of the spiked and non-spiked dataset, we ran a generalized linear mixed model with all the hyperparameters as fixed effect and the models’ loss as random effect. We dropped from the model all the fixed effects that had no significant impact on the prAUC. Thusly, the final model for the absolute difference between class prediction accuracy was(60) (61)

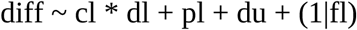

where diff refers to the absolute difference between class prediction accuracy, cl stands for number of convolutional layers, dl for number of dense layers, pl is the pool stride, du dense units as fixed effects and fl the first convolutional layers filter. The residuals followed a Gamma distribution. The final model for the prAUC of the test set was

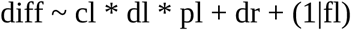

with dr being the dropout rate, and the residuals’ distribution was binomial. We tested the same models in the original complete dataset. Generalized linear mixed models were performed using the package lme4’s function glmer (60) and *p* values were extracted using the function Anova from the R package car (61).

For each type of CNN training we selected the model with the best performance during the testing phase in order to understand the architectures that could offer the best prediction of both classes. We selected models that had a prAUC value on the test set of at least 0.7, and that they did not overfit into predicting one class of DEGs. To do so, we estimated the absolute difference of the accuracy of predicting each class in the dataset. We used the same criteria for all cases.

Once we had selected the model parameters that comprise the architecture which best predicted the DEG class, we evaluated the significance of the result and excluded the possibility that is the result of learning from spurious sequences by permuting the label identity of each gene 100 times and re-train using the same model architecture. We estimated the *p* value by counting the cases that had better performance based on both the prAUC and the class accuracy difference and divided by 100.

All results were visualized in R v4.1.2 (59).

## Results

### The presence of enriched motifs in the up regulated DE gene sequences suggests their potential contribution to environmental response

We searched for motifs enriched along the putative regulatory regions of the up-regulated DEGs in comparison to the non-DEGs to assess the possibility of patterns in the regulatory regions associating with the differential expression status of each gene. We used STREME (49) to search for de novo enriched motifs within the upstream and downstream regions of all genes of interest (Figure 1a). When we searched for enriched motifs only within the sequences of up-regulated DEGs for the reference genotype, *Col-0,* we discovered 9 motifs. In contrast, we identified 14 motifs significantly enriched within the up-regulated DEGs of all 5 accessions, suggesting greater power with more genotypes (*p <* 0.05). The median discovered motif length was 10bp (S3 Table) and the majority of the discovered motifs were significantly enriched in high frequency in our dataset. Specifically, five of the fourteen discovered motifs were present in more than 98% of the DEGs’ regulatory regions tested, even though a few rare ones with less than 1% frequency were also identified (Figure 1b). When we looked at which genotypes each discovered motif was significantly enriched for at least once, the average number of motifs per up-regulated DEG was similar between the genotypes (approximately 6 motifs per gene per genotype; Figure 1c).

**Figure 1:**
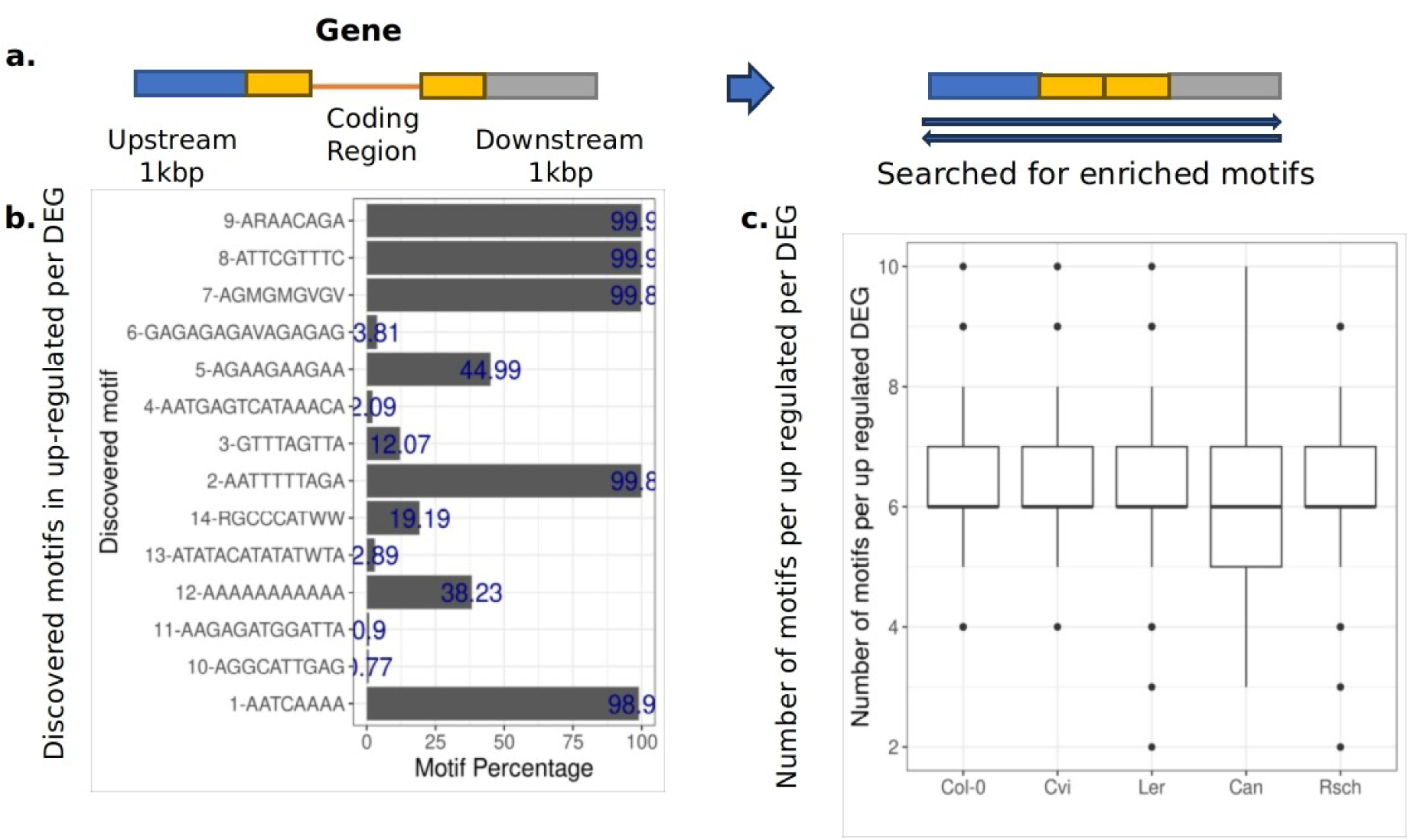
Significantly enriched motifs within the regulatory regions of the up- and down- regulated DEGs. **a.** For each up- or down- regulated gene across all five *A. thaliana* accessions, we extracted the 500 bp upstream (blue block) and downstream regions (grey block). We also extracted within the first 500 bp after the transcriptional starting site and 500 bp before the transcriptional ending (yellow blocks) for a total of 1k bp per gene to include in the motif search. **b,d.** The discovered motifs in the upstream and downstream regions of up- or down- regulated DEGs and how frequent they are in the DEG sequences with STREME. **c,e** The distribution of discovered motifs in the upstream region of up- DEGs or down- DEGs per accession.

There was a similar number of enriched motifs among the down-regulated DEGs. In total 15 motifs were discovered that were significantly enriched in all genotypes (*p* < 0.05; S4 Table). Five of those motifs were present in more than 99% of tested DEG sequences for enrichment (Figure 1d), with an average of 6 discovered motifs per tested DEG sequence (Figure 1e).

When we cross referenced all the motif sequences we identified as enriched in up- and down-regulated DEGs in a database of experimentally defined TFBSs (50), the discovered motifs had a significant overlap with 135 and 143 known TFBS, respectively (*p < 0.05*). The most common TFBS across the discovered motifs were members of the DOF family, with CDF5, DOF1.7 and DOF5.1 binding sites being present in 3 and 4 of the total discovered motifs in the up-regulated and down-regulated DEGs respectively. The most common discovered sequences overlapping with the members of the DOF family were ARAACAGA in up-regulated and CAAAAAAAA in down-regulated DEGs. Members of the DOF family of transcription factors have been characterized in being involved in cold response(62). Moreover, *PISTILLATA* (PI), member of the MADS-box factors family MIKC, and the C2H2 zinc finger factor SGR5 binding sites, were present in 3 and 4 of the discovered motifs in the up -regulated DEGs, respectively. There was an overlap of the most common TFBS across the discovered motifs of the up-regulated DEGs and the down-regulated DEGs (Table 1). Therefore, within the promoter regions of the up- and down-regulated DEGs, there are motifs present that could be potentially informative for classifying the gene expression.

**Table 1:** The top known transcription factor binding sites (TFBS) that significantly overlap with the enriched discovered motifs along the upstream and downstream regions of up and down- regulated regions of up and down DEGs.

| TFBS | Number of discovered motifs | Gene | Family |
| --- | --- | --- | --- |
| MA1159.1 | 4 | SGR5 | C2H2 zinc finger factors (class) |
| MA1268<br>.1 | 3 | CDF5 | DOF |
| MA0556<br>.1 | 3 | DOF1.7 | DOF |
| MA0558<br>.1 | 3 | FLC | MIKC |
| MA0559<br>.1 | 3 | (PISTILLATA) PI | MIKC |
| MA0940<br>.1 | 3 | AP1 | MADS box factors<br>(AP1) |
| MA1085<br>.1 | 3 | WRKY40 | WRKY (class) |
| MA1281<br>.1 | 3 | DOF5.1 | DOF |
| MA1367<br>.1 | 3 | AT1G76870 | Trihelix |
| MA1380<br>.1 | 3 | TCX6 | CPP |
| MA1182<br>.1 | 5 | RVE8 | Myb-related |
| MA1184<br>.1 | 5 | RVE1 | Myb-related |
| MA1380<br>.1 | 5 | TCX6 | CPP |
| MA1277<br>.1 | 4 | DOF1.7 | DOF |
| MA1281<br>.1 | 4 | DOF5.1 | DOF |
| MA1268<br>.1 | 4 | CDF5 | DOF |
| MA1278<br>.1 | 4 | DOF3.4 | DOF |
| MA1279<br>.1 | 4 | DOF1.5 | DOF |
| MA1190<br>.1 | 4 | RVE5 | Myb-related |
| MA0933<br>.1 | 4 | AHL20 | HMGA factors |
| MA1267<br>.1 | 4 | DOF5.8 | DOF |

### Known transcription factor binding sites within *Col-0* do not accurately predict the expression response to cold using random forests

Given the many known and documented transcription factor binding site motifs that exist for different transcription factors (many of which are known to be involved in cold response) and given that we found TFBS motifs in the cold DEGs, we asked if these TFBS could predict genome-wide expression response to cold. We used a dataset of known TFBS in putative promoters 1kbp upstream of genes in the *A. thaliana* accession *Col-0* (52) to test whether they can predict DEGs using random forests classification trees (S1 Figure). For this analysis, we also split the genes based on up-regulated and down-regulated and compared each group to all non-DEGs. Even though both random forests models correctly predicted the gene regulation status for approximately 80% of the genes, the prAUC of each of those models was low. The probability AUC-ROC (prAUC) for predicting up-regulated DEGs was 0.5013 (permutation based p = 0.17) and the prAUC for predicting down-regulated DEGs was 0.499 (permutation based *p* = 0.22). Therefore, RF models based on known TFBS involved in temperature related reactions were not able to predict DEGs with high accuracy within this dataset. This, taken together with the various discovered motifs within the upstream and downstream regions of the genes, indicates that a method that incorporates more genetic information within the code might improve the prediction of regulatory status better within specific conditions.

### Predicting gene expression regulation based on regulatory regions using naive methods is possible

We next investigated whether a currently commonly used machine learning method could learn *cis* regulatory motifs simply from sequence and DE data alone. If *de novo* motifs can be identified in DE genes, even though known TFBS were not good predictors of expression in the RF model, we might obtain better predictions learning *de novo* important motifs. To this end, we trained convolutional neural networks (CNNs) that incorporate the upstream and downstream genetic sequence as input layer via one hot encoding (Figure 2a; see Material and Methods). We trained 1,344 models with different combinations of the hidden layer’s parameters to find the model architecture that best predicts the genes DE (S2 Table). We trained the models using the information from all 5 accessions that had available pseudogenomes to predict up- or down-regulated DEGs separately.

**Figure 2:**
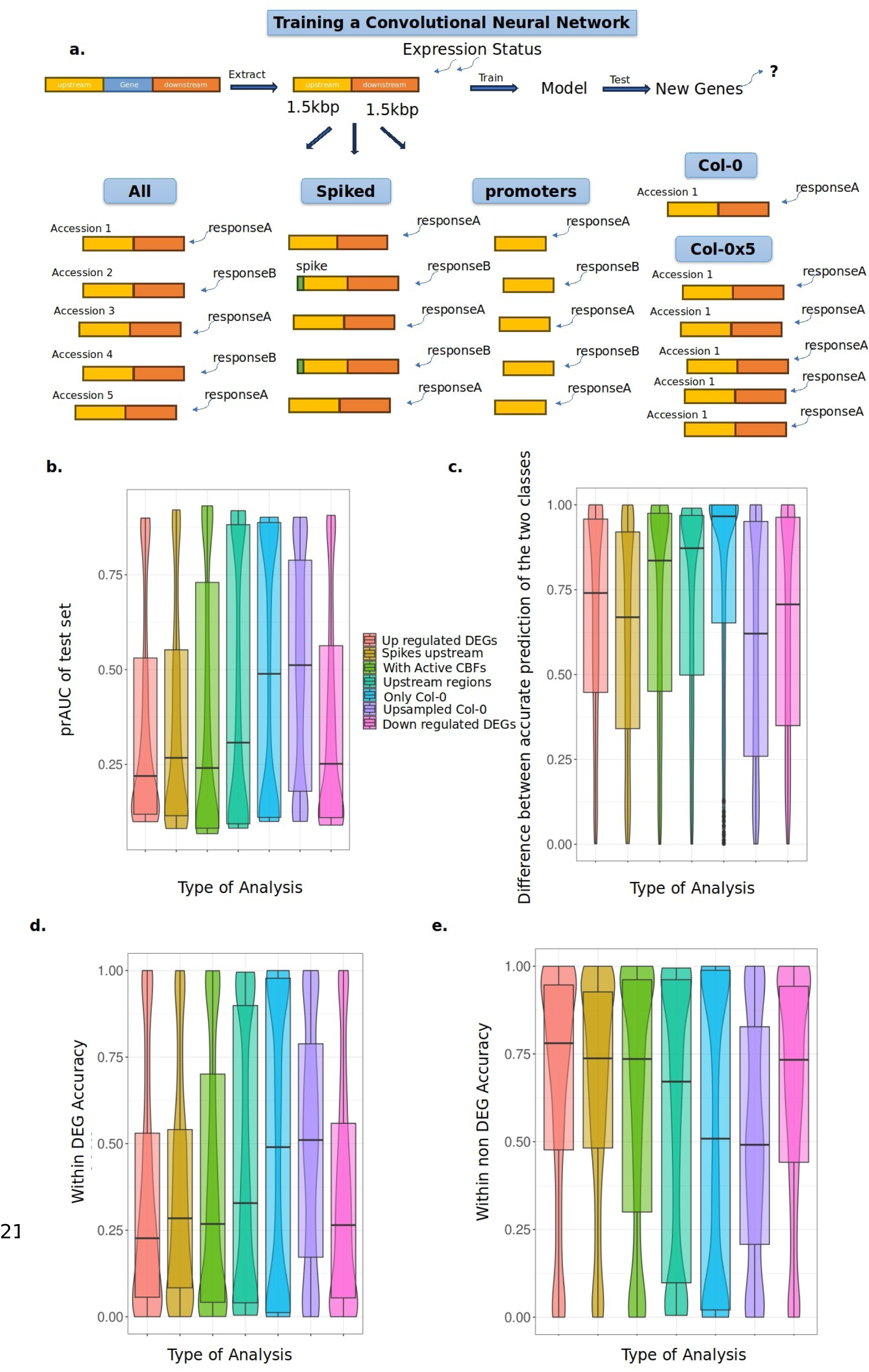
Training Convolutional Neural Networks to predict up-regulated or down-regulated DEGs between specific environments. a. Description of the different type of input data used to train CNNs to predict up- or down-regulated DEGs vs non DEGs. The analysis “up-regulated DEGs” included the five *A.thaliana* accessions and both upstream and downstream regions for up-regulated DEGs. The analysis “spiked upstream” included training on up-regulated DEGs with the first 5bp being replaced consistently with the same sequence. “With active CBFs” refers to the training of models using the two accessions with all *3 CBF* genes active. The analysis “upstream regions” included only the upstream regions of the up-regulated DEGs as input. “Only *Col-0”* was used to train models, as well as when it was five times oversampled (“Col-0×5”). Finally, the results under the category “down regulated DEGs” show the outcome of training in all **b.** prAUC values of the test set in the different analysis. **c.** The difference in accurately predicting the up/down-regulated DEGs to the non DEGs in all different analysis. **d.** The within class prediction accuracy of down- and up- regulated DEGs in the test set in all analysis. The within class prediction accuracy was estimated as the proportion of correctly predicted genes of this class over the total number of genes in this class in the test set. **e**. The within non-DEGs prediction accuracy of the test set in all analysis.

The tested CNN model architectures have had a wide variety of accuracy in predicting the DEG of each gene (S5 Table). We extracted the prAUC of each model during training and moreover, we estimated the prAUC of the test set for each trained CNN model. The median and mean prAUC values during training for predicting up-regulated DEGs vs non-DEGs were 0.5013 and 0.5017, respectively, indicating that many tuning parameter settings did not lead to substantial learning. We also estimated the median within class accuracy in the test set as 0.776 and 0.227 for non DEGs and up-regulated DEGs, respectively. Note that the test set is not over sampled, but it is the original ratio which is mostly non-DEGs. Overall, the median prAUC in the test set of the models was 0.229, with a mean of 0.354 (S2 Figure). We identified 15.6% of the models with prAUC in the test set values above 0.80.

To gain insight into whether tuning parameters had any consistent effects on model learning, we used a generalized linear mixed on the prAUC of the test set using the first layer’s filters as random effect and the number of convolutional and dense layers, with pool stride and dropout rate as fixed effects. Both the number of convolutional and dense layers had a significant negative impact on the prAUC of the test set, with *p* = 0.000754 and *p* = 3.9708e-16, respectively, suggesting more complex models were more poorly trained. Additionally, the pool stride (*p* = 2.918e-10), which indicates the number of base pairs that are considered together in each filter, and the dropout rate, or how many model nodes are used each time to predict, had a significant negative impact of the prAUC of the test set. However, none of the interactions between the fixed effects were significant, suggesting little importance of specific combinations of the tuning parameters on learning.

The trained models for predicting down-regulated DEG or non-DEGs (S3 Figure) were very similar with the results observed for the up-regulated DEGs vs non-DEGs. The median prAUC of the models during training was 0.5009 and the mean prAUC was 0.5012. The median prAUC of the test set for the down-regulated DEGs vs non DEGs was 0.253 and the mean 0.363 (S6 Table). The same hyperparameters as for predicting up-regulated DEGs were identified to have a significant negative effect with *p-values* of 1.511e-07, 0.0168, 4.068e-08 and 0.0485 for number of convolutional layers, number of dense layers, pool stride and dropout respectively.

We identified the best models that could predict the up-regulated DEGs. For that purpose, we selected a model that had prAUC in the test set above 0.7. Moreover, in order to understand the models’ potential to predict each class separately we estimated for each class a prediction accuracy as the number of correct predictions of the class divided by the total number of the genes in the class tested. So, after this prAUC cutoff, we picked a model whose prediction accuracy difference was the lowest. The best model for predicting up regulated DEGs had a prAUC in the test set of 0.701, and prediction accuracy in the test set was 0.368 for up regulated and 0.693 for non DEG (Table 2). The class prediction accuracy difference was 0.325. The best model for predicting up-regulated DEGs had 2 convolutional layers and 2 dense layers, a dropout rate of 0.25 and pool stride 4. Using the same criteria, we were able to identify the best model in predicting the down-regulated DEGs. The prAUC of the test set for this model is 0.7128 and the difference in the class prediction accuracy is 0.319, with per class accuracy being 0.365 for down-regulated DEGs and 0.665 for non-DEGs. The model had 3 convolutional and 2 dense layers, pool stride of 8 and dropout rate of 0.25.

**Table 2:** The best model for each training performed are given. For each type of analysis run (Figure 2), we selected the best performing model based on the prAUC value of the test set and the the difference in accurately predicting the DEGs vs non DEGs. For each model, the model parameters, the training, validation and testing prAUC, loss and accuracy of predicting both DEG classes is given.

|  | Up regulate<br>d DEGs | Down<br>regulat<br>ed<br>DEGs | Spiked<br>up<br>regulat<br>ed<br>DEGs | Upstrea<br>m<br>regions<br>of up<br>DEGs | Up<br>regulat<br>ed<br>DEGs of<br>only | Up<br>regulat<br>ed<br>DEGs of<br>upsamp | Genot<br>ypes<br>with<br>all *CBF*<br>as |
| --- | --- | --- | --- | --- | --- | --- | --- |

|  |  |  |  |  | *Col-0* | *led Col-0* | DEGs. |
| --- | --- | --- | --- | --- | --- | --- | --- |
| 1st conv. filters | 128 | 64 | 128 | 64 | 128 | 128 | 128 |
| 2nd conv. filters | 128 | 64 | None | 128 | None | 128 | 128 |
| 3rd conv filters | None | 128 | None | 128 | None | None | 64 |
| Conv width | 4 | 4 | 4 | 4 | 4 | 8 | 8 |
| Pool width | 4 | 4 | 8 | 4 | 4 | 4 | 4 |
| Pool stride | 4 | 8 | 4 | 4 | 8 | 4 | 8 |
| Dropout | 0.25 | 0.25 | 0.25 | 0.1 | 0.1 | 0.1 | 0.25 |
| 1<sup>st</sup> dense layer units | 64 | 128 | 128 | 128 | 128 | 128 | 64 |
| 2<sup>nd</sup> dense units | 128 | None | 128 | None | 128 | 128 | None |
| Number of conv. layers | 2 | 3 | 1 | 3 | 1 | 2 | 3 |
| Number of dense layers | 3 | 2 | 3 | 2 | 3 | 3 | 2 |
| Learning rate | 0.0001 | 0.0001 | 0.0001 | 0.0001 | 0.0001 | 0.0001 | 0.0001 |
| Loss | 0.692 | 0.691 | 0.691 | 0.691 | 0.691 | 0.692 | 0.691 |
| Accuracy | 0.621 | 0.682 | 0.673 | 0.666 | 0.692 | 0.636 | 0.672 |
| Accuracy of DEGs | 0.368 | 0.365 | 0.480 | 0.339 | 0.347 | 0.368 | 0.341 |
| Accuracy of non DEGs | 0.693 | 0.684 | 0.665 | 0.671 | 0.697 | 0.681 | 0.681 |
| prAUC test set | 0.701 | 0.712 | 0.704 | 0.701 | 0.7207<br>444 | 0.703 | 0.711 |
| prAUC training | 0.499 | 0.502 | 0.522 | 0.499 | 0.4971<br>713 | 0.505 | 0.499 |
| prAUC of validation set | 0.501 | 0.502 | 0.578 | 0.509 | 0.4851<br>478 | 0.517 | 0.506 |
| Loss of the validation set | 0.693 | 0.693 | 0.692 | 0.693 | 0.6933<br>79 | 0.693 | 0.693 |
| Epoch of best model | 37 | 50 | 49 | 29 | 49 | 48 | 49 |
| Difference in accuracy of DEG classes | 0.325 | 0.319 | 0.185 | 0.332 | 0.3495<br>906 | 0.312 | 0.340 |

We assessed whether these models performed better than randomness by permuting the input labels and re-train the selected model 100 times (S4 Figure), so as to understand to what degree the input labels are used for tuning by the CNN architecture. For each permutation we estimated the prAUC of the test set and the prediction accuracy between the two predicted classes using the un-permuted test set. Then, we compared the values of the original trained model against the distribution of the metrics estimated with the permuted labels. The significantly higher prAUC for the up-regulated DEGs (*p* < 2.2e-16) and down-regulated DEGs (*p* < 2.2e-16) than the permuted dataset indicate that our picked models perform better than randomly picking genes.

### Number of discovered motifs can impact the correct prediction of up-regulated DEGs

We investigated the profile of the genes that are accurately predicted within the up regulated DEGs by the selected model with the highest achieved performance. Out of the 4,564 up DEGs across all five genotypes in the test set, 592 were accurately predicted. The correctly predicted genes had a significantly higher (wilcoxon rank sum test *p* = 0.018) average number of discovered motifs in their regulatory regions (median of 12 motifs) by the STREME analysis than the genes that were incorrectly predicted (mean of 10; S5a Figure). Interestingly, when we repeated this analysis for the down-regulated DEGs, we did not detect a different accumulation of TFBS within the correctly predicted genes. Both the correctly and wrongly predicted genes had an median of 11 motifs per gene and the accumulation of the TFBS upstream and downstream of the regions was not significantly different (wilcox sum rank test *p* = 0.9446; S5b Figure). Moreover, the gene expression level of the genes that were correctly predicted vs not correctly predicted was not significantly different (based on Kolmogorov Smirnov test p > 0.05) for both down-regulated DEGs and up-regulated DEGs.

Up until this point, the discovered motif diversity, the CNN training results, and even the best model’s behavior was very similar for up-regulated DEGs versus down-regulated DEGs. For the rest of the analyses, we focused on training CNN models for predicting up regulated DEGs versus non DEG, as we do not expect that altering the input of the CNNs training will alter the observed pattern between the two regulatory groups.

### CNNs imperfectly identify a simple artificial signal within the regulatory regions

Because many tuning parameter values resulted in models that learned little (training prAUC near 0.5), we sought to assess potential sources of constraints on these models. We investigated whether the tested model architectures can learn a simple motif within the sequences that perfectly predicts DEGs, a motif we refer to as a “spike”. If these spiked models are perfectly able to predict DEGs in this setting, it suggests that the limitations we observed with real data arise from a lack of such good, simple predictors in the sequence. If models have limited learning in this setting, it suggests that the model architecture is limited in its ability to find signals in the data were using, perhaps due to the complexity and abundance of sequence data (2,000 bp per gene).

To test these hypotheses, we included a spike sequence of 5bp in the place of the first 5bp of all up-regulated DEGs (Figure 2a). The spike’s sequence (AAGGG) does not overlap with the sequence of any of the discovered enriched sequences with the up DEGs. The expectation was that the accuracy of the models would increase as they have a perfect sequence to use for predicting the up-DEG group. During training the prAUC values had median of 0.5018 and mean of 0.5046. The median prAUC in the test set of the spiked models was 0.36, with a mean of 0.266, which was on average higher than when trained without the spikes (S7 Table). The distributions of the prAUC values in the test set between the spiked and non-spiked models were significantly different (Kolmogorov Smirnoff test, *p* < 2.2e-16; Figure 2b). Generally, the spiked models had fewer overfitted models than the non-spiked models for both up-regulated and non-DEG classes (Figure 2c; S6 Figure). In total 188 spiked models with prAUC values of the test set above 0.8 were detected and 561 models with prAUC values of the test set below 0.2, and thus, in total, 55.7% of the models tested are overfitted to one of the two classes. In contrast, among the non-spiked models, 61.5% of the models are overfitted.

When we used the same criteria to pick the best performing models for the spiked data (permutation *p <* 2.2e-16) as for the non-spiked data, we identify a model that predicts the two classes better than the non-spiked best model. prAUC of the test set (0.704) is only marginally higher by 0.03. In contrast, the difference in the prediction accuracy of the two classes in the test set is 0.185, which is 1.75 times lower than the best model predicting non-spiked up-regulated DEGs. This improvement is due to the higher accurate prediction of the up-regulated DEGs (0.48) in the spiked best model than the non-spiked best model (0.36).

There is no significant correlation between the prAUC values in the test set of the spiked and non-spiked models under the same values of tuning parameters (*p*=0.96, ρ =0.0012), indicating that the inclusion of the spikes did not improve models with good tuning parameters for the real data, but instead were best captured with distinct tuning parameters (Figure 3). When we further investigated the different hyperparameters’ impact on the prediction accuracy of each class, we noticed that the spiked models with 1 convolutional layer had higher accuracy and prAUC in the test set and simultaneously smaller difference between the accurate prediction of both classes than the models with 2 and 3 convolutional layers. (Figure 3a). The fact that the models with less convolutional layers performing better that with more of them was also true for the spiked models. A generalized mixed model including the number of convolutional layers, dense layers, pool stride and dense units as fixed effects indicated that the number of convolutional layers had a significant effect on the difference of the prediction accuracy of the two classes (*p* = 8.091e-10). Fewer layers were predicting more accurately, as the model’s estimates were negative. This observed pattern is not as extreme as for the non-spiked models (*p* = 0.01023; Figure 3a). Other parameters that were found to have a significant negative impact on the spiked models’ difference in the prediction accuracy of the two classes were the number of dense layers ( *p* = 9.036e-05), the pool stride (*p* = 0.002137), the first dense layer’s units (*p=*0.048), as well as the interaction of the number of convolutional and dense layers (*p* = 0.00983). All these factors, except for the interaction of the number of dense and convolutional layers, had a significant effect on the difference between the accuracy of the two classes for the non-spiked models. Specifically, the number of dense layers, the pool stride, and the number of convolutional layers had *p* values of 6.191e-06 and 0.0102 respectively (Figure 3b-c). The estimates for all parameters were negative in both spiked and non-spiked data, which means that generally models with fewer layers were able to predict with higher per class accuracy in the test set. This result suggests that although different specific sets of tuning parameters for good for spiked data versus the non-spiked, the general effect of tuning parameters, and specifically model complexity, was consistent. Therefore, a consistent signal within the up-regulated DEGs can improve the model performance, indicating that the regulatory complexity among genotypes is quite high, that we lack information in the sequence data for prediction, or that the DEG response is highly stochastic.

**Figure 3:**
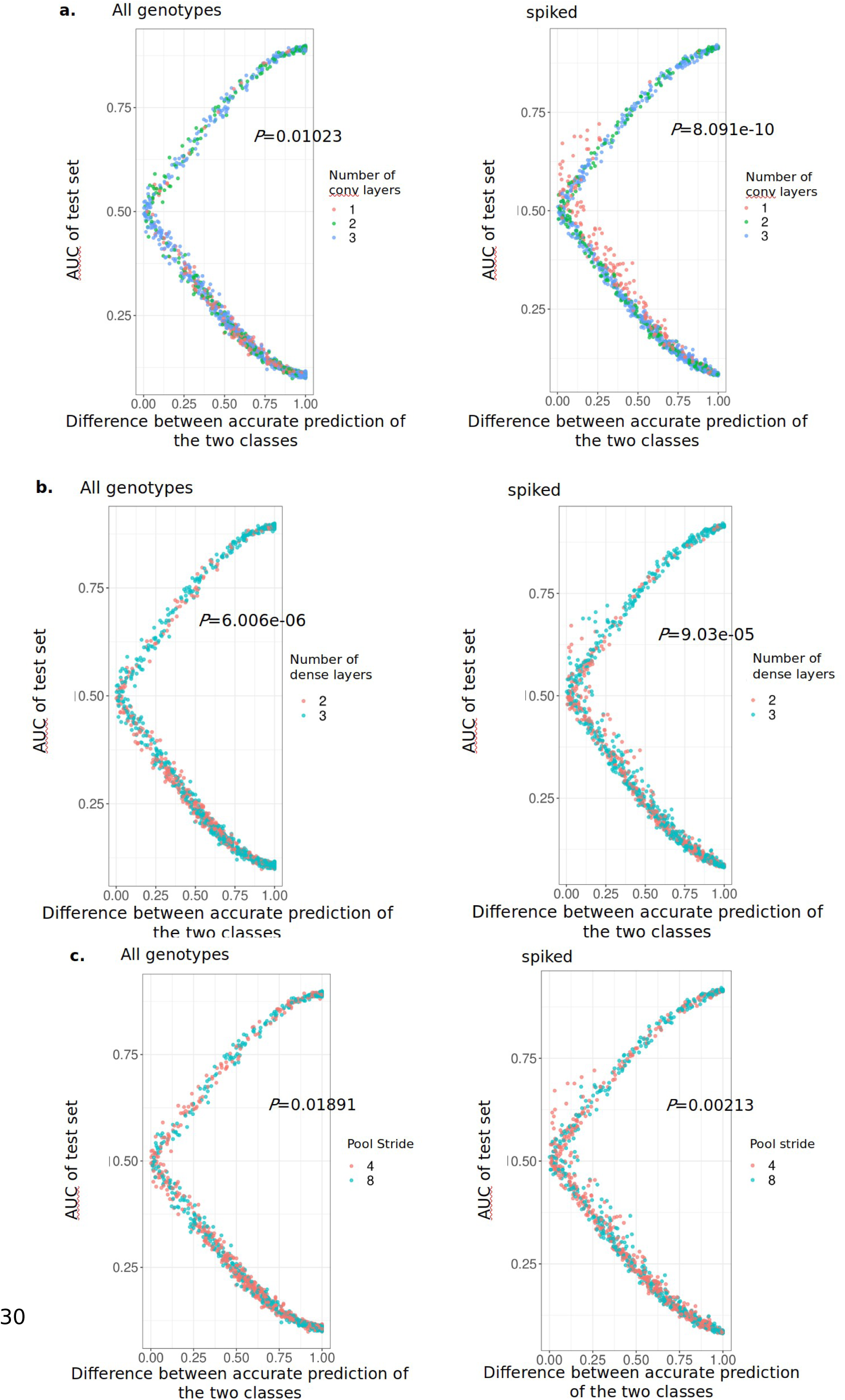
Simple spiked models can accurately predict an introduced spike of 5bp at the beginning of the up-regulated DEGs promoter region. Comparison of the prAUC in the test set and the difference between the accurate prediction of the two classes for non- spiked (left) and spiked (right) input data. Comparisons are done for a. number of convolutional layers, b. number of dense layers and c. pool stride. The p values stand for the significance level of the KS test when comparing the two distributions shown within each plot.

### Predicting only among genotypes sharing alleles for major *trans* eQTL does not improve the prediction of the up-regulated DEGs

Since the results with the sequence spike indicate some data complexity limited our CNN predictions, we investigated whether we could simplify the prediction task by homogenizing known large-effect *trans* mutations (CBFs). These transcription factors show variation in functionality among genotypes, and in the other natural genotypes tested at least some copies are non-functional. We approached this analysis in two ways. First, we trained models predicting up-regulated DEGs vs non DEGs in *Rsch, Cvi* and *Ler*, which do not have intronic SNP differences between them in the two accessions that have all three major well-studied cold-response *CBF* transcription factors, including *CBF2*. Then, we trained models to predict up-regulated vs non DEGs on *Can* and *Cvi,* which have differential expression of all 3 *CBFs* during the experiment.

The median and mean prAUC during training on the set with no SNP differences, was 0.5001 and 0.5 respectively, while the prAUC of the test set had mean value of 0.47 and median 0.095 (S7 Figure; S8 Table). This is also reflected in the percentage of model architectures, which have been overfitted for predicting the upregulated DEGs; 52.5%. of the models in contrast to the 45.5% overfitted in predicting the non DEGs, with only 1.7% of the models not showing any sign of overfitting. The best model trained on only the three accessions with similar copies of *CBF* functional had prAUC in the test set of 0.705 and difference in predicting accuracy of the two DEG groups of 0.47. Similarly, the median prAUC during training on the set with differentially expressed all 3 *CBFs*, was 0.5014, while the prAUC of the test set had median value of 0.268 (S7 Figure; S8 Table). The difference in the accurate prediction of the up-regulated DEGs and non DEGs within the test set has a median of 0.835 and mean 0.701, which is on average higher than the difference in prediction accuracy of the models applied to all genotypes. The best model had prAUC in the test set of 0.7 and difference in predicting accuracy of the two DEG groups of 0.31. Overall, those models did not perform better than the original models including all the *A. thaliana* accessions. This suggests that the lack of a major *trans* acting expression variant did not hinder predictability in our bigger set of diverse genotypes here.

### Including both upstream and downstream regions improves the prediction of up-regulated DEG class

As often the *cis* elements are mostly identified in the upstream regions (64), we tested whether including the downstream coding region has limited the prediction accuracy, perhaps by incorporating noise. Noise might be introduced by incorporating long sequences that do not usually have a lot of TFBS, such as the downstream regions. Therefore, we retrained all the models by including only the 2kbp upstream regions for all 5 accessions, compared to the 1 kb up and 1kb downstream of the coding region in the approach presented above (Figure 2a). The median prAUC of the training set was 0.5 and the median prAUC (0.307) in the test set of the models trained only using the 2kb upstream sequences was slightly higher than including both the 1kb upstream and 1kb downstream regions, by 0.048 (S8 Figure; S9 Table). The distribution of the prAUC results in the test set of the two analysis was again significantly different (KS test *p* < 2.2e-16). The within class prediction accuracy of up-regulated DEGs (0.671) in the test set was higher than for non-DEGs (0.328). Moreover, 29.8% of the models had prAUC values in the test set above 0.8. The same pattern was observed for models that had low prAUC in the test set. Approximately 41% of the models had prAUC in the test set below 0.2, indicating overfitting to predicting all genes as up-regulated DEGs, to a total of 71.1% of the models overfitting to either class. In comparison, including both upstream and downstream regions leads to overfitting of 61.5% of the models. (Figure 2c, Figure 2d). Therefore, excluding the downstream regions from the analysis leads to more cases when the models get fixed in recognizing only one class.

The best model in predicting up-regulated DEGs and non DEGs trained only on putative promoter regions had 3 convolutional layers, 3 dense layers and a dropout of 0.1 (Table 2). The prAUC in the test set was 0.7004 and the difference in accurately predicting up-regulated DEGs and no DEGs is 0.332, which is a similar performance to the best model learning using both upstream and downstream regions (Table 2). Its performance is better than models with the same architecture trained on random input (*p* < 2.2e-16). Therefore, including only the 2kb upstream regulatory regions of the genes during training does not significantly alter the predictive ability compared to including 1kb upstream and 1kb downstream.

### Genetic diversity does not improve accuracy of differentially expressed genes

Genetic diversity is an important dimension of natural populations. Incorporating multiple genotypes in the STREME analysis has had a positive impact into discovering more enriched motifs. If we consider the introduced spike in the upstream region of up-regulated DEGs as a large effect mutations due to its 100% frequency within the group, we also showed that the majority of the information that the models learn is based on small effect mutations. Small effect mutations can be better identified when incorporating more genetic diversity into any kind of analysis, as the less frequent variants are easier to identify. We finally compared how effective CNNs were when including multiple genotypes in the training set, by training all the different model architectures including only the most commonly studied *A. thaliana* accession, *Col-0* (Figure 2a). We repeated this twice, once that had the original number of genes in *Col-0* and once that we up sampled the *Col-0* genes enough times to have the same size of training set as for the complete set of the genotypes. In both cases, we oversampled the up regulated DEGs to a 1:1 ratio with the no DEGs. The training of the non-oversampled and oversampled sets wielded median prAUC values of 0.5004 and 0.501 respectively.

When compared to the models that have been trained on the *Col-0* dataset only, we see that the median AUC of the test set is higher, as it is at 0.488, for the multiple genotypes model (Figure 2b, S9 Figure; S10 Table). The accuracy was significantly impacted by the number of replicates, as the models trained on the up sampled *Col-0* set had median prAUC of the test 0.508 (KS test *p* = 8.8e-16; S11 Table; S10 Figure). During training, both datasets had a high proportion of overfitted models. Within the up sampled dataset, 50.3% of the prAUC values in the test set were either above 0.8 or below 0.2. Training on the up-sampled dataset of *Col-0* had fewer overfitted models than training on the non-oversampled dataset of *Col-0,* in which 77.4% of the models were overfitting in either direction. The better performance of the oversampled dataset of *Col-0* is due to the more appropriate size of the dataset; the original *Col-0* dataset has only 34,002 data points when the up-regulated DEGs are oversampled to 1:1 ratio (65,66). Based on this observation, below we compare only the models trained on the oversampled dataset of *Col-0* to the models trained on all five accessions.

The models trained on the *Col-0* dataset have fewer overfitted models (50.3%) than the models trained on all five accessions (61.5%). The best model in predicting both up-regulated and non-DEGs on the *Col-0* dataset has similar architecture and performance statistics as the best model trained on all five accessions. The prAUC value in the test set trained on the oversampled *Col-0* was 0.7034 and the difference in the accurate prediction between the two classes was 0.3122. It performed better than when trained on a permuted dataset (*p* < 2.2e-16). Both the prAUC in the test set (0.7008) and the difference in accurate prediction of both classes (0.3252) indicate that training on all five accessions was performing not as well as the model trained on the *Col-0* dataset only. Therefore, including more accessions may either marginally increase the noise and not improve the accuracy of the models by a lot, or not be variable enough to capture more contrasts between alleles rather than genes.

Finally, because the models of cold response had only modest success, we tried to test if the CNN models could accurately predict DEGs in a simpler transcriptomic change (1,282 DEGs), based on a single transcription factor knockout known to control anthocyanin biosynthesis. However, even these CNN predictions had modest success (Supplemental Material), highlighting the challenge in predicting transcriptomic dynamics from proximate DNA sequences.

## Discussion

We used the genomic sequence of upstream and downstream regions of *Arabidopsis thaliana* genes to identify motifs related to the differential expression in cold and control conditions and also to predict the gene expression response to an abiotic stressor. Many *cis* regulatory elements have been documented (13,38,63) and they are often involved in gene evolution and adaptation to novel environments (4,67). Here, within both the up-regulated and down-regulated DEGs we found motifs of variable length that are enriched within them in comparison to the non-DEGs. The discovered motifs in *cis* regulatory regions significantly overlap with known transcription factor binding sites that are involved in the regulation of gene expression under cold stress (42,62,68). Knowledge of specific transcription factor binding sites can aid the prediction of gene expression under different types of abiotic and biotic stress (26,33,69,70). The five accessions used in this study are regulated by differential function of *CBF* and *ZAT12* regulons, leading to their variable acclimation capacity to cold (42). The three *CBF* genes control the acclimation to cold conditions and reaction to freezing stress (58,71). It has been shown that genetic diversity within the gene and its network is connected to adaptation to different temperatures and latitudes (9,45,72). We identified a frequent overlap with known TFBS, which are known to be involved in controlling responses to cold or freezing stress such as the RVEs (73), members of the DOF family of transcription factors (58,62,74,75) and FLC (76). We also identified frequent binding sites for members of the DREAM complex, which are involved in repressing growth in response to DNA damage (77), as well as SGR5, which is differentially spliced under heat stress (78). This indicates the possibility to predict gene expression regulation based not only on factors with biological meaning, but also with genetic diversity, as the incorporation of more accessions than *Col-0*, yielded more discovered motif sequences.

However, as often the available information to the researcher is only DNA and mRNA sequence (and not experimentally verified transcription factor binding sites), we investigated the possibility of using the genetic sequence to predict gene expression. The need for this was also evident by trying to predict up- or down-regulated DEGs using random forest analysis. The random forest used known presence or absence of specific TFBS upstream and downstream of the genes to predict DEGs. However, neither for up-regulated DEGs nor for down-regulated DEGs, the prediction accuracy was better than chance. In *A. thaliana,* using known transcription factor binding sites has not yielded very good prediction of differential gene expression (26), while the most enriched *cis* regulatory elements did not also predict the most differentially expressed genes in maize (32). Training CNNs, with the 2kbp upstream and downstream regions of the genes wielded better prediction accuracy. The model that had both high accuracy and had the least difference in predicting accuracy per class had prAUC in the test set of 0.7. This value is slightly lower than what has been reported in similar studies in other species, which identified models with prAUC between 75% to 85% (55,79). In our case, higher prAUC values showed signs of overfitting; non DEGs, which comprise more than 80% of the dataset were then almost exclusively correctly predicted, inflating the accuracy and prAUC. Surprisingly, the inclusion of both upstream and downstream regulatory regions did not improve the prediction accuracy of DEGs, as in maize (80). This could be attributed to genomic differences between the species; for instance downstream regulatory regions have differences in nucleotide composition between maize and *A. thaliana* (81). Additionally, maize has a different genomic composition than *A. thaliana,* such as higher transcription element density, with 85% in contrast to 21% in *A. thaliana.* Authors have suggested the expanded genome of maize could enhance the landscape and availability of *cis* regulatory adaptations (82). Factors such as this can explain the difficulty of successfully transferring machine learning protocols generated from one species to another.

Phenotypic responses to environment (plasticity) can be adaptive, which are ultimately fixed by populations in the specific environment (85). Phenotypic plasticity can also be genetically variable (i.e. GxE), with gene expression being an important example. Our hypothesis was that incorporating more genotypes in this study would improve the prediction accuracy as more genetic variation in *cis* sequences and expression would be incorporated during training. Incorporating genetic variability between species has been successfully used to train models with simple architecture to predict gene network relationships (86). Within maize, when more than one genotype was used for predicting gene expression, then the results were more accurate (33). We did not observe the same pattern here; in fact an oversampled dataset of *Col-0* upstream and downstream regions to the same size as the dataset including the multiple genotypes, yielded a model with higher prAUC and more comparable prediction of the two DEG classes. We do not believe that this is simply an issue of power due to the dataset being on the smaller side of what is required for machine learning. A potential contributing factor could be that the *cis* -regulatory genetic variation might not be of large enough frequency between genotypes but is stronger between genes. Additionally, the highest prAUC in the test set was achieved when the accessions with differentially expressed all 3 *CBF* genes were only included during training, suggesting *trans* effects in high frequency could limit our predictions across multiple genotypes. This pattern was confirmed when a high frequency motif was incorporated in the upstream regions of the up-regulated DEGs; the spiked dataset could be used to train models with higher accuracy. Within *A. thaliana,* the CBF/DREBs genes of the cold signaling pathway have been linked with cold adaptation within Europe (44,87) as the major eQTLs controlling acclimation. Taken together with the fact that more than half of the enriched discovered motifs within the up- and down-regulated DEGs are present in less than 50% of the genes across all genotypes and thus not very frequent, it indicates that the large number of *cis* regulatory variants across the genome in low frequency can decrease the accuracy in prediction of gene expression differences, in contrast to the presence of a potential large effect eQTL controlling the response to cold. The putative promoter region likely captures the many *cis*-regulatory elements, which have been related to large impacts on gene expression divergence and polygenic selection but are often each individually mutations of small effect (63,88). Incorporating a spike within the upstream region, which represents a single factor of large effect, supports this hypothesis, as in overall the spike improved the prediction accuracy of the models. Our findings are consistent with machine learning predictions of protein folding, where models that accurately predict variation among genes perform worse on predicting effects of individual amino acid substitutions (89), a task that was better suited to a distinct prediction approach (90). During training, the models were able to detect the spike and increase the accuracy of the up-regulated DEGs. Therefore, even though including multiple accessions could slightly increase the accuracy of predicting GxE, care has to be taken to which accessions are selected for the training.

One explanation for the remaining variation in expression unpredicted by CNNs in this study is that these models overlook much of the biological complexity driving gene expression response to environments. Genes are expressed as a part of complicated regulatory networks that *cis* regulatory elements only partially capture. We have currently ignored epigenetic states and the sequence of the coding region. In a sorghum species, it has been more successful to predict DEGs as response to cold as models incorporated information about the coding region sequence and methylation information (79). Alongside specific transcription factor binding sites, their positions on the genome and the chromatin state have been useful to predict the transcription factor’s target genes, with information that is also transferable with high accuracy between datasets (84). Prediction accuracy of gene expression differences and the impact of *cis* variants were improved when large up- and downstream regions, as well as when open methylated regions were included in the analysis (32,33). Trained bimodal CNNs based on chromatin state and DNA sequence for different cell types can differentially predict the transcription factor binding specificity, which performs better than neural networks based only on DNA sequence information (83). During training, therefore, using only *cis* regulatory regions to predict gene expression response to environment within *A. thaliana* may be of limited accuracy without incorporating additional data types. However, when that biological information is missing and it is challenging to acquire, as would be the case in many non-model systems, the accuracy of imperfect CNN models might still be of use to the evolutionary researcher.

Interpretation of what might have been having an impact on the training could be difficult to understand. Even models with the best performance do not necessarily have the best interpretable structure of layers (91). Looking into the statistics of all the different models trained during each grid search can give insights into this. First, the distribution of the prAUC during testing can give clear indications of how easy it is to overfit to the model, and therefore how complex this training task can be. Despite these limitations, using machine learning methods can enable researchers to draw predictions and conclusions about reactions to specific environments within species, albeit perhaps not CNN models. We believe that further development of other models such as language models, which can use less biological information to predict gene expression responses, would be useful for systems with less information about their (molecular) biology than the model species.

## Acknowledgments

The Pennsylvania State University ACI cluster and the HPC Ramses of University of Cologne have provided us with computational support. We would like to thank S. Mahony and H. Dittberner for providing valuable feedback in running convolutional neural network analysis and M.G. Stetter, T.S. Winkler, A. Singh and J. Floret for feedback on the manuscript.

## Data Accessibility

All scripts are available at the github repositories: https://github.com/mtakou/cnns_DEGpromoters and https://github.com/em-bellis/CNN_GxE.

