## Supplemental Figures for "Predicting gene expression responses to environment in *Arabidopsis thaliana* using natural variation in DNA sequence"

**Supplemental Material**

We tried to assess if the CNN models can accurately predict DEGs by the promoter regions, when there is one functional change in a gene network. This represents a biological signal similar to our spiked dataset, and also contrasting it between genetic backgrounds. For this reason, we used DEGs between *Col-0* and a knock out line of the TFBS *tt8*, which have documented differences in the regulation of glycosylation (47). We used the up-regulated DEGs, which were more numerous than the down-regulated DEGs and we encoded *Col-0* for the input to the model. The median prAUC value of the training sets was 0.5 and the median prAUC in the test set was 0.967. The distribution of the prAUC values during grid search was bimodal, with the majority of the values indicating overfitting (S11a Figure). However, we identified the best model in predicting with similar accuracy both down- and non DEGs, with values of 0.263 and 0.741 respectively, that had a prAUC in the training set of 0.745 (S11 Figure). Therefore, including more accessions may either marginally increase the noise and not improve the accuracy of the models by a lot, or not be variable enough to capture more contrasts between alleles rather than genes.

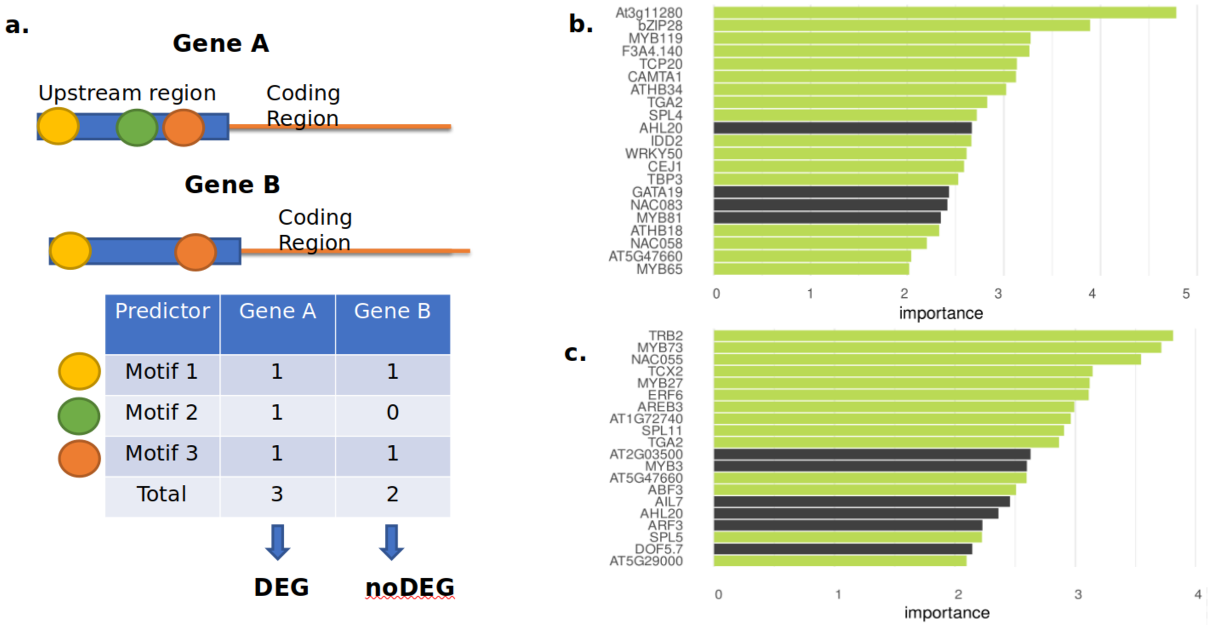


S1 Figure**:** Known transcription factor binding sites of *Col-0* poorly predict the expression response to cold, with PRAUC = 0.503. **a.** We used a database of known TFBS upstream of *Col-0* to predict whether a gene is DEG or non DEG. Each TFBS was scored as being present (1) or not present (0) in each feature. **b.** The random forest features with the highest relative importance within the up regulated DEGs within *Col-0.* **c.** *T*he random forest features with the highest relative importance for predicting down-regulated DEGs within *Col-0*. In green and black it is marked if a gene's relative importance is significant or not significant, respectively, based on *p* values less than 0.05 The *p* value for each feature was estimated based on 100 permutations of the dataset.

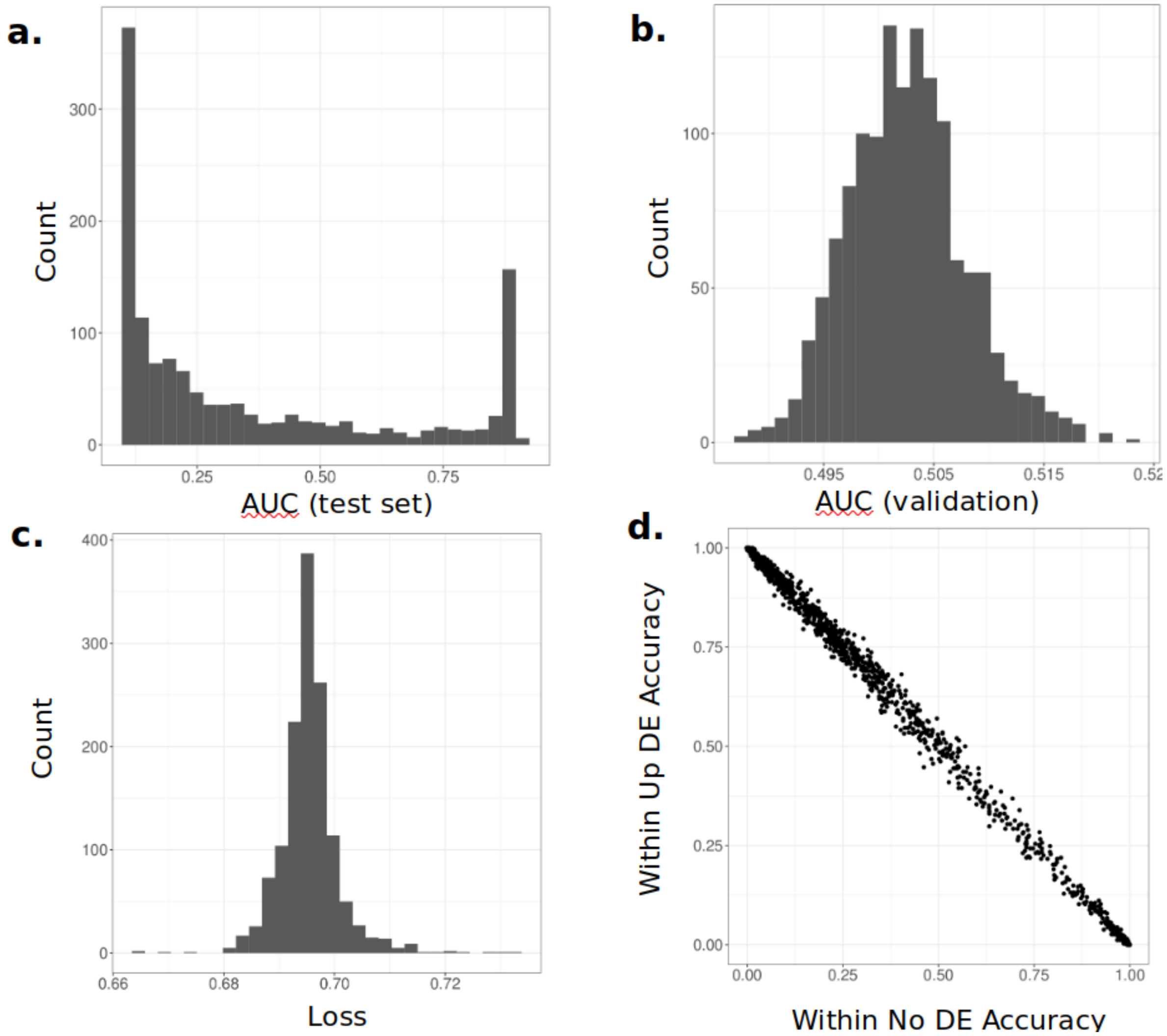


S2 Figure: Summary statistics of all models trained during the grid search for predicting up-regulated DEGs vs non DEGs. The models were evaluated by estimating the a. prAUC of the test set, b. the prAUC of the validation set during training, c. the loss of each model during training and the d. accuracy of correctly predicting each class in the test set, The distributions of summary statistics for all tested CNN architectures are given.

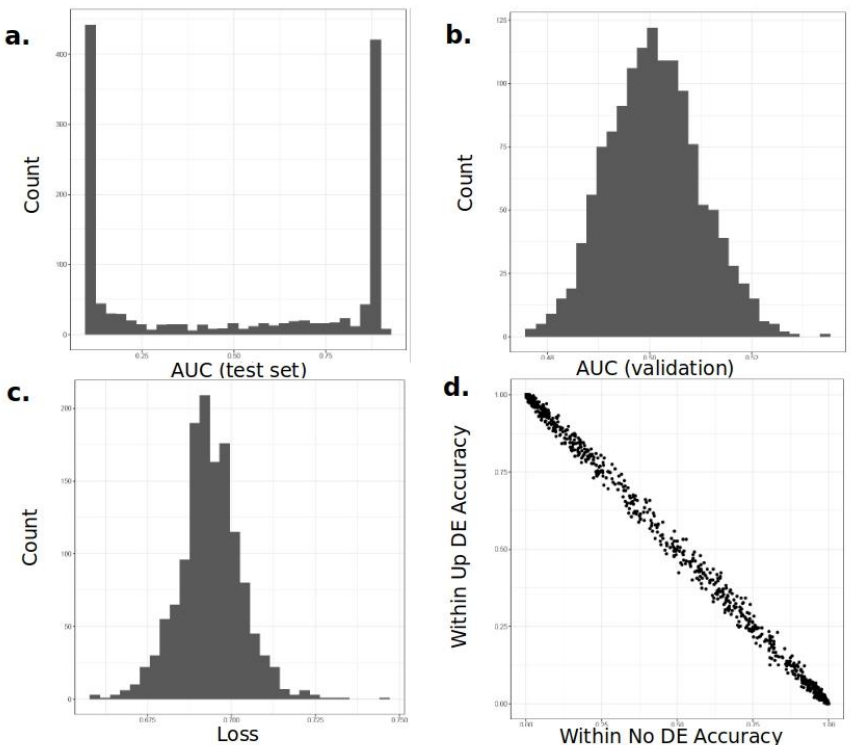


S3 Figure: Summary statistics of all models trained during the grid search for predicting down-regulated DEGs vs non DEGs. The models were evaluated by estimating the a. prAUC of the test set, b. the prAUC of the validation set during training, c. the loss of each model during training and the d. accuracy of correctly predicting each class in the test set, The distributions of summary statistics for all tested CNN architectures are given.

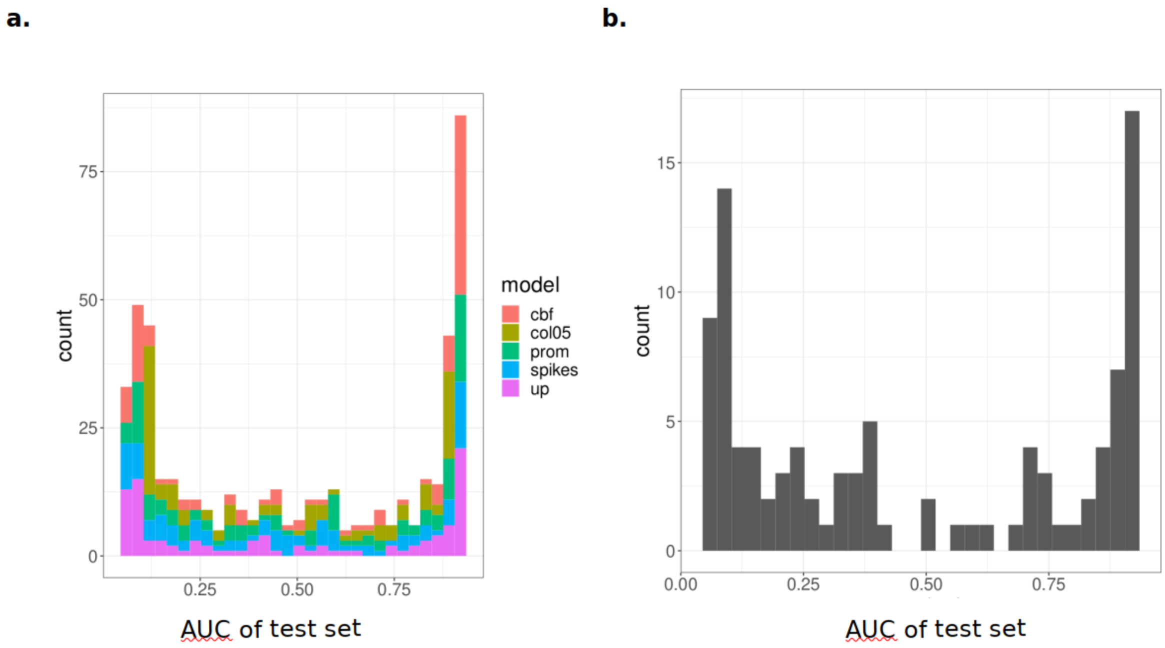


S4 Figure: Distribution of the prAUC in the test set for the permuted datasets of a) up-regulated DEGs (up) and b) down-regulated prAUC,.For the up-regulated DEGs we permuted the labels for all the subsequent analysis, specifically for when only the 2 accessions with all 3 CBFs are active (cbf), for up-sampled *Col-0* dataset (col05), for only the upstream regulatory regions included in the analysis (prom) and so when a spike as positive control was incorporated (spikes).

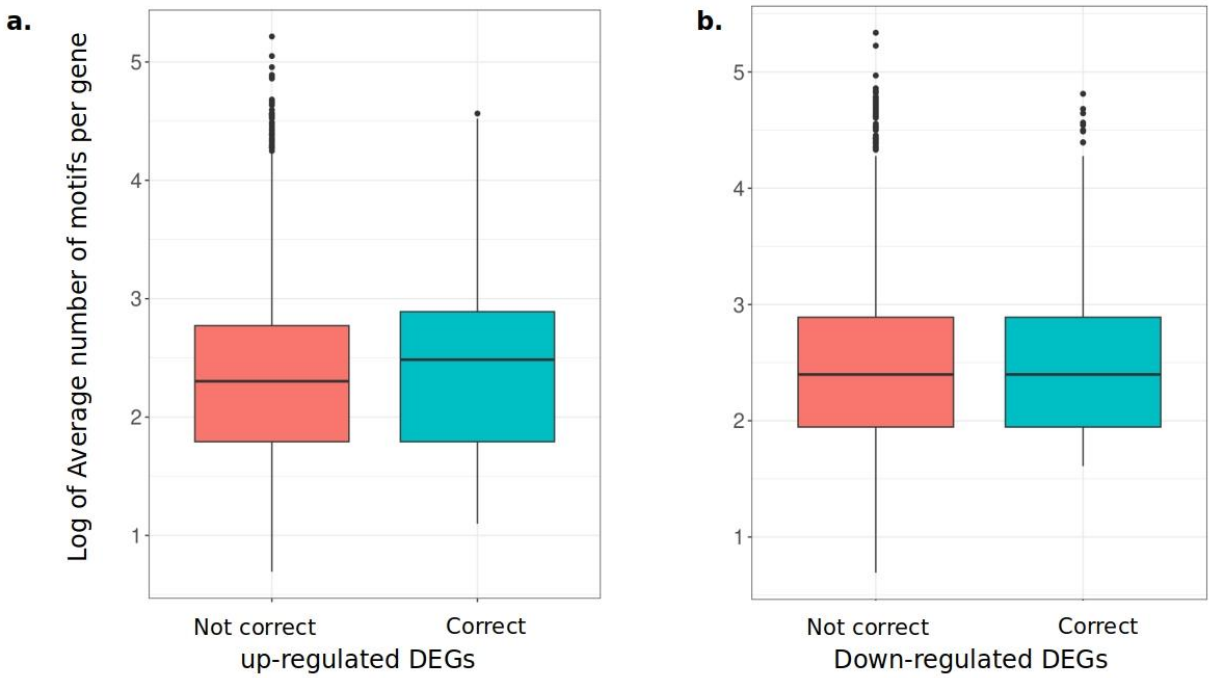


S5 Figure: The average number of motifs per gene for the correctly predicted and not correctly predicted DEGs by the best model predicting a. up-regulated DEGs or b. down regulated DEGs.

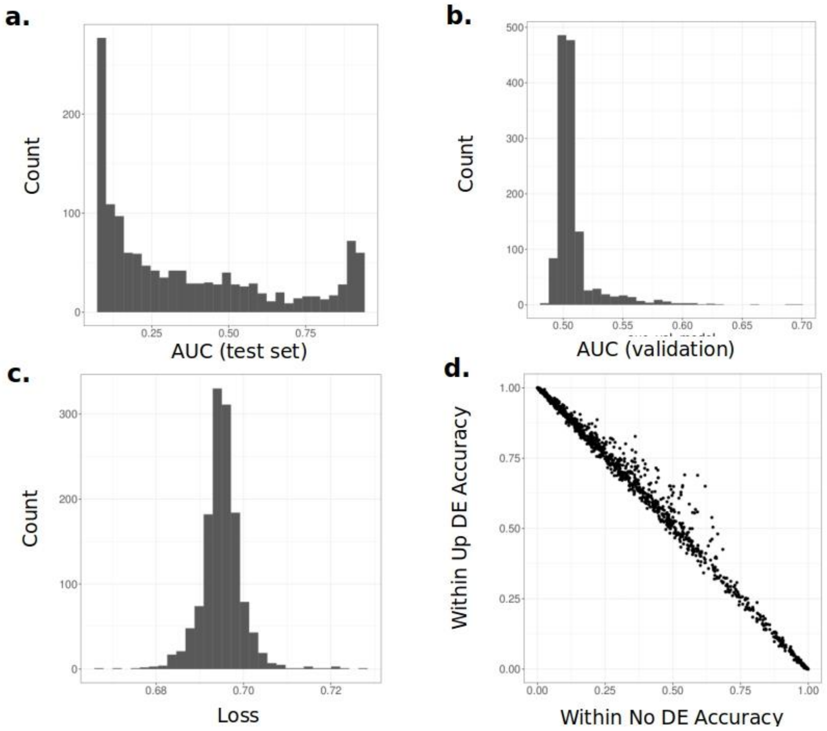


S6 Figure: Summary statistics of all models trained during the grid search for predicting up-regulated DEGs vs no DEGs, while including a spiked region at the start of the up-regulated genes' putative promoters. The models were evaluated by estimating the a. prAUC of the test set, b. the prAUC of the validation set during training, c. the loss of each model during training and the d. accuracy of correctly predicting each class.

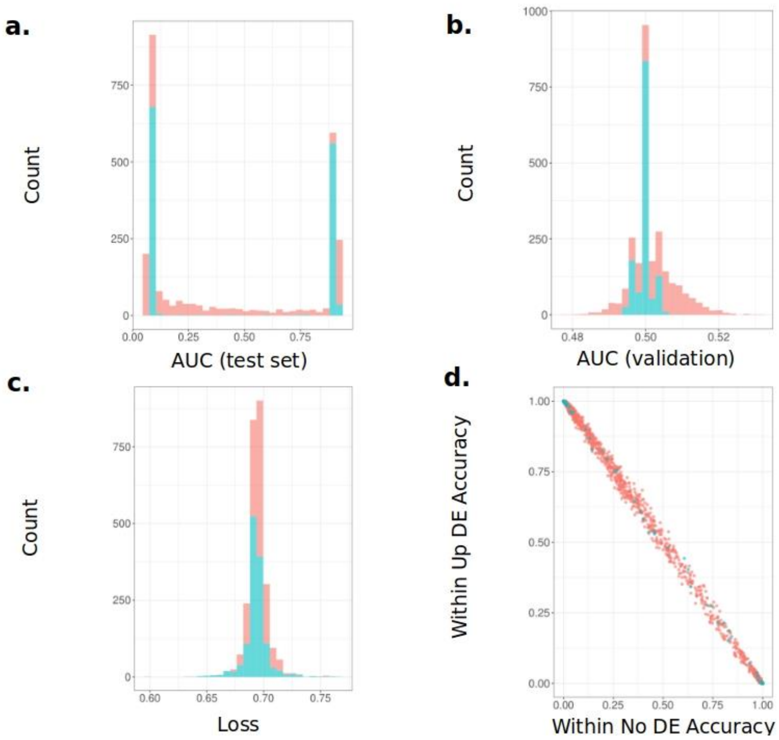


S7 Figure: Summary statistics of all models trained during the grid search for predicting up-regulated DEGs vs non DEGs, while using the two accessions with all 3 *CBF* genes differentially expressed between the two treatments (red) and all 3 *CBF* genes having no SNP differences between the accessions. The models were evaluated by estimating the a. prAUC of the test set, b. the prAUC of the validation set during training, c. the loss of each model during training and the d. accuracy of correctly predicting each class, The distributions of summary statistics for all tested CNN architectures are given.

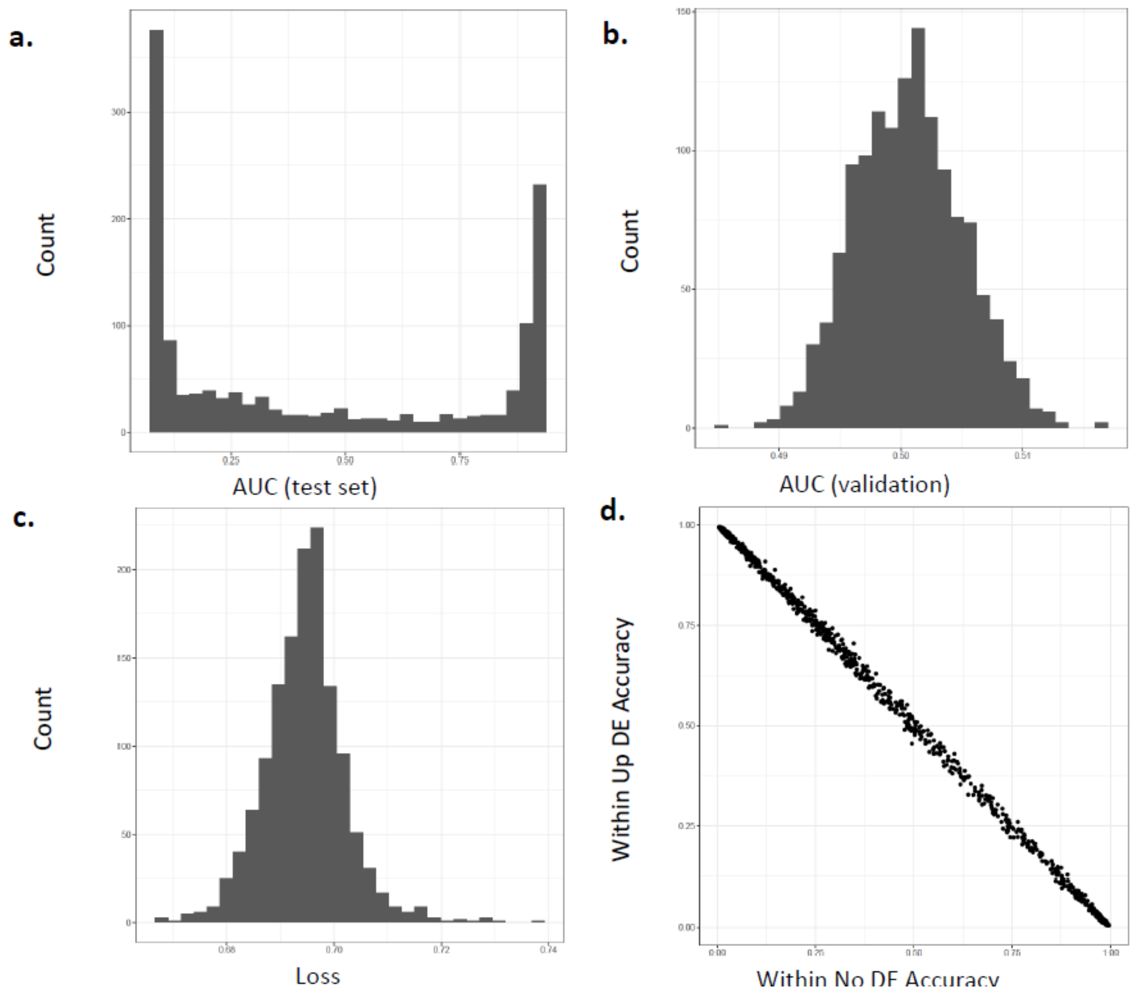


S8 Figure: Summary statistics of all models trained during the grid search for predicting up-regulated DEGs vs non DEGs, while including only up-regulated genes' putative promoters. The models were evaluated by estimating the a. prAUC of the test set, b. the prAUC of the validation set during training, c. the loss of each model during training and the d. accuracy of correctly predicting each class, The distributions of summary statistics for all tested CNN architectures are given.

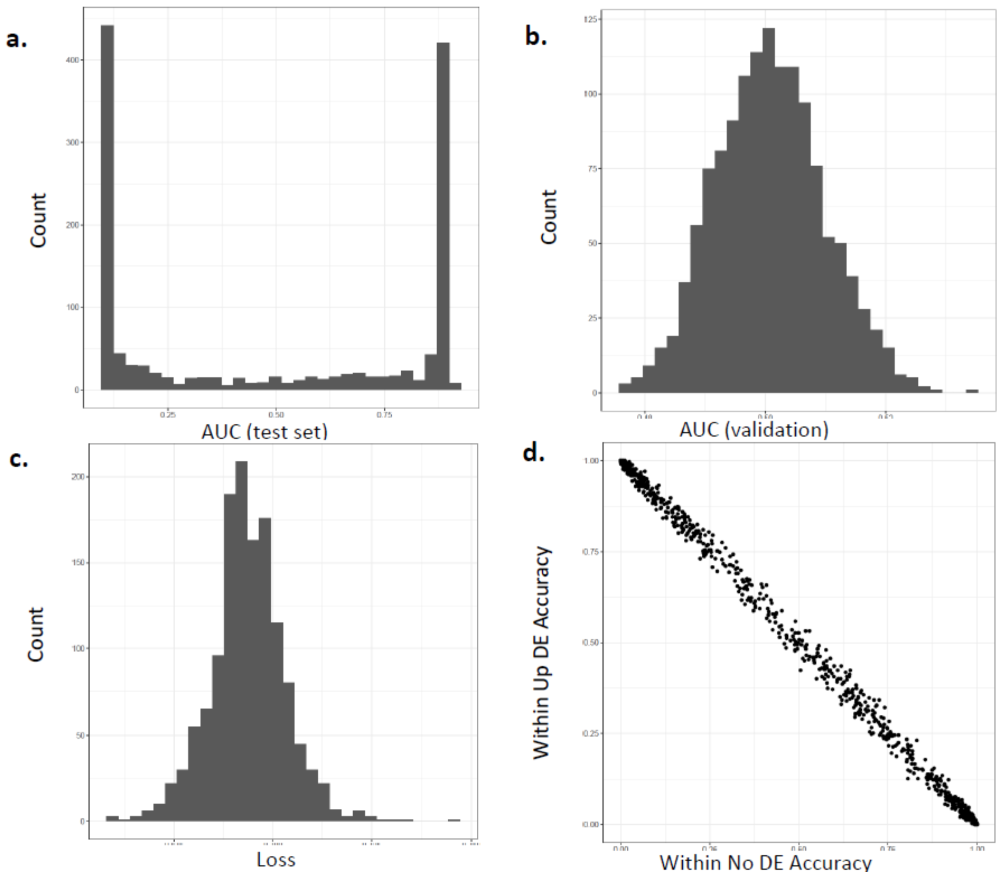


S9 Figure: Summary statistics of all models trained during the grid search for predicting up-regulated DEGs vs no DEGs, while including only *Col-0*.The models were evaluated by estimating the a. prAUC of the test set, b. the prAUC of the validation set during training, c. the loss of each model during training and the d. accuracy of correctly predicting each class, The distributions of summary statistics for all tested CNN architectures are given.

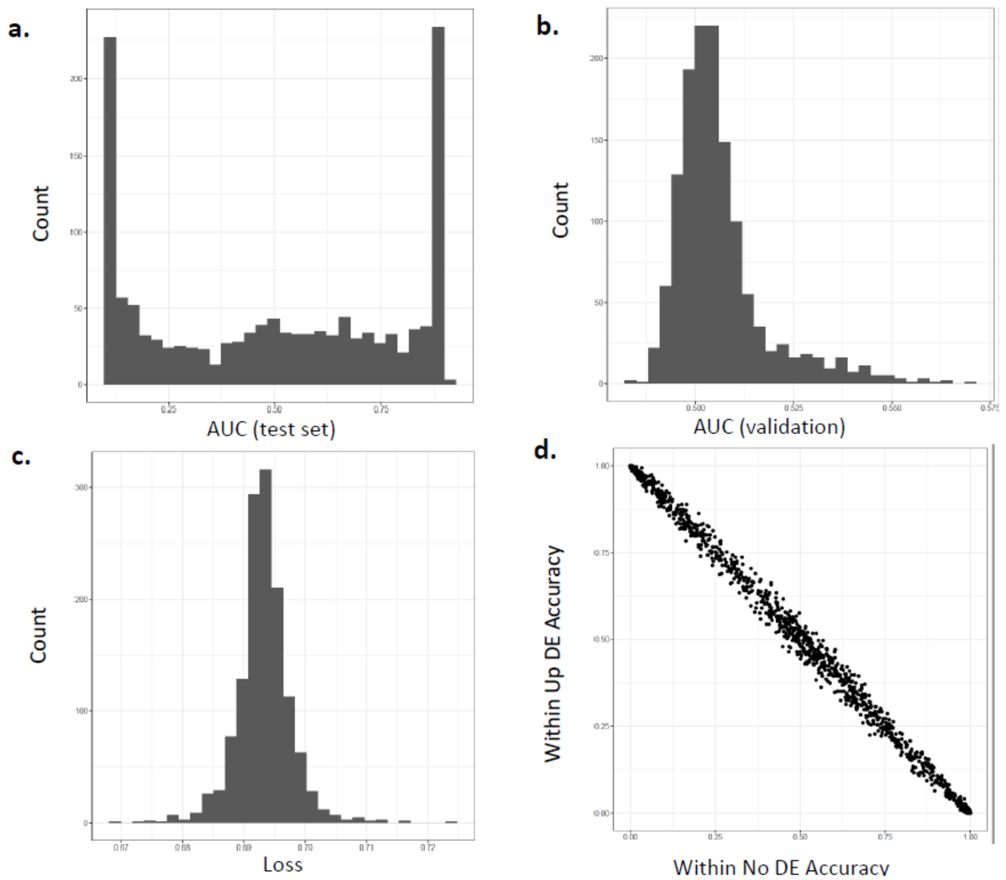


S10 Figure: Summary statistics of all models trained during the grid search for predicting up-regulated DEGs vs no DEGs, while including only an up sampled set of *Col-0*. The models were evaluated by estimating the a. prAUC of the test set, b. the prAUC of the validation set during training, c. the loss of each model during training and the d. accuracy of correctly predicting each class, The distributions of summary statistics for all tested CNN architectures are given.

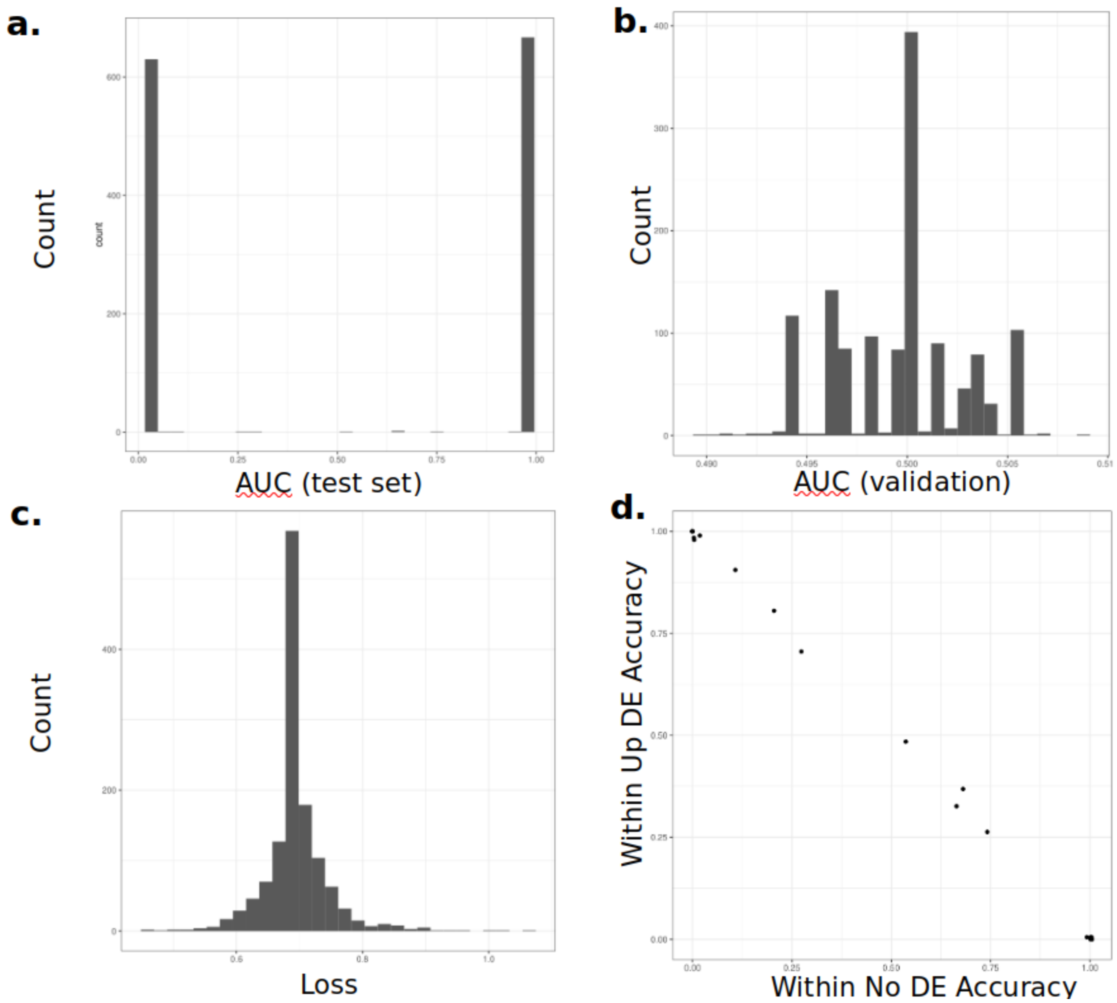

S11 Figure: Summary statistics of all models trained during the grid search for predicting up-regulated DEGs vs no DEGs in an up sampled set of a *tt8* knock out dataset. The models were evaluated by estimating the a. prAUC of the test set, b. the prAUC of the validation set during training, c. the loss of each model during training and the d. accuracy of correctly predicting each class, The distributions of summary statistics for all tested CNN architectures are given.
